# Actin Assembly on Biomimetic Lipid-coated and Functionalized Bead Surfaces

**DOI:** 10.64898/2026.11.11.765950

**Authors:** Olivia L. Mooren, Patrick McConnell, Anshuman Jaysingh, John A. Cooper

**Affiliations:** Department of Biochemistry and Molecular Biophysics, Washington University School of Medicine, St. Louis, MO

## Abstract

Dynamic assembly of actin filaments drives cell protrusion and other aspects of cell motility. Reconstitution of actin assembly from purified components is a powerful approach for understanding the molecular mechanisms that underlie these processes. Purified preparations of actin, actin-binding proteins, and regulators of actin-binding proteins enable a broad spectrum of reconstitution experiments. Much of this actin filament assembly occurs at the plasma membrane, with exquisite temporal and spatial regulation. We developed a biomimetic reconstitution system using purified lipids or glutathione on the surface of functionalized beads as models for actin assembly at the plasma membrane. The system incorporates two critical pathways for regulation of actin assembly – nucleation of actin polymerization by Arp2/3 complex and activation of heterodimeric actin capping protein (CP). Arp2/3 complex and CP are both essentially inactive in the basal resting state of the cytoplasm. The system incorporates membrane-bound activators for Arp2/3 complex, such as WASP and WAVE family proteins, and for CP, such as CARMILs and other CPI-motif proteins. Membrane-bound components are coupled to beads either through lipids, which allow lateral mobility, or through glutathione groups attached to the bead surface via flexible linkers. Fluorescence microscopy of labeled components, combined with quantitative image analysis, is used to assess the combined effects of the regulators and thereby test molecular mechanisms for regulation of actin assembly. Here, we describe protocols for preparing functionalized beads, coupling regulators to the bead surface, performing actin assembly reactions, and imaging and quantifying the resulting actin networks.

**Summary:** To investigate actin assembly at membranes, we developed reconstitution systems using beads as membrane mimics. We targeted regulators to lipid-coated or glutathione-functionalized beads. Actin polymerization was induced with purified Arp2/3 complex, capping protein (CP), and their regulators. Actin network assembly was quantified by fluorescence microscopy.

## Introduction

Understanding the molecular mechanisms by which actin filaments interact with the plasma membrane is a fundamental issue in cell motility ^1, 2^. Barbed ends of actin filaments are oriented toward the membrane. The ends can be stably attached as structural elements to effect contractile actions, or they can be free to polymerize or depolymerize, powering protrusion or retraction of the membrane.

Biochemical and genetic approaches have identified a host of actin-binding proteins that contribute to actin filament assembly and disassembly, along with many components, both proteins and lipids, that control the activity of the actin-binding proteins ^3^. Modern methods of protein purification, from native and recombinant sources, have enabled the use of increasingly complex mixtures of regulators and components in biochemical reconstitution experiments ^4, 5^.

Studies of moving cells, including assessment of individual components, have shown that actin monomers are readily available in the cytoplasm and that their polymerization into filaments can be stimulated and controlled by several factors that act at the plasma membrane ^6^. Nucleation of actin filament assembly by Arp2/3 complex is a major mechanism for stimulation of actin polymerization at membranes ^7, 8^. Arp2/3 complex activity can be regulated in several ways, including by nucleation-promoting factors (NPFs) that bind directly to Arp2/3 complex and promote a conformational change from the resting inactive state to the active one. Arp2/3-mediated assembly requires the presence of a barbed-end capping factor, notably the heterodimeric alpha/beta capping protein (CP) ^9^. CP is present at high concentrations in cytoplasm but held in an inactive state by the protein V-1, which binds tightly to the actin-binding site of CP, acting as a simple steric blocker ^10^. CP can be activated by CPI-motif proteins, such as CARMILs, which bind directly to a separate site on CP and induce a conformational change that promotes the dissociation of V-1, thus activating CP ^11^.

Biochemical reconstitution systems have provided powerful evidence about the possible mechanisms by which multiple components combine to stimulate and regulate actin polymerization. A seminal study by Carlier and colleagues demonstrated that actin assembly could drive motility and be induced by a defined set of proteins including Arp2/3 complex, CP, and ADF/cofilin ^12^. That study used an NPF on the surface of bacteria to recruit and activate Arp2/3 complex. Actin assembly and bacterial movement depended on the concentration of Arp2/3 complex, CP, and ADF/cofilin, which were found to be essential components. The concentration-dependence displayed a bell-shaped curve, providing insights into the molecular mechanisms involved. Other components, including VASP, profilin, and alpha-actinin, were not essential, but they provided additional layers of regulation.

Inspired by those findings, the field of actin assembly and actin-based motility has developed increasingly complex reconstitution systems that have provided new insights into molecular and cellular mechanisms ^13–15^. Reconstitution of actin assembly inside cell-sized liposomes has provided a model for assembly of actin filaments and the actin cortex ^16, 17^. We have developed a bead-based system that incorporates a mimic for the plasma membrane by coating beads with lipids. We added NPFs for Arp2/3 complex and activators of CP to the surface using non-covalent mechanisms that allow the components to move laterally as they would in a membrane. As an alternative, we attached GST-tagged proteins to glutathione-functionalized polystyrene beads. While these attachments do not move laterally, the amount of each component on the bead can be determined with confidence, and they provide a control for the lateral mobility of lipid-targeted components. To these coated beads, we added Arp2/3 complex, CP, V-1, and profilin-actin to induce actin polymerization. This system faithfully recreates key features of membrane-associated actin assembly, and we use it to test molecular mechanisms. Our system incorporates fluorescence imaging of labeled components, allowing us to localize the components relative to the bead and the actin filament network and to quantify their levels.

Our methods allow for easy fine-tuning of multiple components to readily observe the effects on actin organization and growth. Specific attachment of membrane-localized proteins, by GST/glutathione or Ni/His couplings, allows for greater control over how much is at the bead surface. Overall, we believe that utilizing the outlined methods to observe actin network growth in the presence of specific proteins at a membrane surface will be beneficial to any in the field of actin regulation.

### Protocol

This protocol provides detailed methods for the biomimetic bead assays reported in our original papers ^18, 19^. We modified well-established protocols ^4, 12, 15, 20, 21^ so that we could fine-tune the activity of the regulator proteins of interest. We also highlight details to consider when setting up these experiments.

## 1. Buffer Preparation

### 1.1. Bead assay buffer

Prepare bead assay buffer with 20 mM HEPES, 100 mM KCl, 1 mM MgCl_2_, 1 mM EGTA, 1 mM TCEP, pH 7.0. Sterile-filter, aliquot, and store this base buffer at -20 °C. Add 1 mM ATP fresh on the day of the assay.

1.1.1. For bead actin polymerization microscopy assays, make a 10x stock of 2% methyl cellulose (400 cP) and 25 mg/mL bovine serum albumin (BSA) to reduce Brownian motion and non-specific sticking to the coverslip. Add this stock to the final bead reaction mixture.

1.1.2. For bead actin polymerization microscopy assays, make a 200 mM beta- mercaptoethanol (βME, 10x stock) to reduce photobleaching. Add this to the final bead reaction mixture.

### 1.2. MOPS buffer

Prepare liposome buffer and protein-coupling buffer with 20.0 mM MOPS, 100 mM KCl, 1.0 mM NaN_3_, 1.0 mM TCEP, pH 7.2. Use this buffer for making liposomes, coating silica beads with lipid, and coupling of proteins to the bead surface.

### 1.3. 2x arrest mixture

Prepare 20 µM phalloidin, 20 µM latrunculin B, 40 mM βME, 5 mg/mL BSA, and 0.4% methyl cellulose in bead assay buffer without ATP. Use this mixture to stop actin polymerization before imaging or antibody labeling (steps 4.4 and 4.5).

CAUTION: Follow all proper chemical safety instructions for using chemicals and wear appropriate personal protective equipment (PPE) while making buffers. Sodium azide is acutely toxic. βME is toxic and malodorous; handle it in a fume hood. Phalloidin and latrunculin B are toxic; wear gloves when handling them.

## 2. Glutathione-functionalized bead preparation

To better control how much protein is attached to each bead, we use glutathione-functionalized polystyrene beads.

NOTE: In this assay, GST-VVCA is the portion of N-WASP that activates Arp2/3 complex; VVCA is tandem Verprolin homology, Central hydrophobic, Acidic. Verprolin homology domain is also known as WH2 for Wasp Homology 2 ^22, 23^. GST-CBR115 is a portion of CARMIL1 that regulates CP activity; it includes both Capping Protein Interacting (CPI) and CARMIL Specific Interacting (CSI) domains ^24, 25^.

2.1. Vortex glutathione-functionalized beads with a diameter of 2 µm for 30 s to resuspend beads.

2.1.1. Sonicate bead suspension in a water bath for 10 min.
2.1.2. Centrifuge for 10 min at 2,400 x g.
2.1.3. Wash once with bead assay buffer (no ATP).

2.2. Resuspend beads in assay buffer (no ATP) containing the following concentrations of GST fusion proteins: 2 µM GST-VVCA, 0-2 µM GST-CBR115, 0-2 µM GST. GST alone is used as a placeholder when the concentration of GST-CBR115 is less than 2 µM, so that the total concentration of GST is 4 µM in all samples during the coating process.

2.3. Incubate beads with GST fusion proteins for 2 h on ice, with frequent agitation to maintain the beads in suspension.

2.4. Centrifuge and wash beads once in assay buffer (no ATP) containing 1% BSA, then again in assay buffer (no ATP) containing 0.1% BSA, and finally resuspend in assay buffer containing 1 mM ATP and 0.1% BSA.

2.5. To determine the number of beads in each protein-coupled sample, measure the OD600. Generate a standard curve of OD600 values for bead suspensions of known concentrations. Adjust the volume for each sample so that the same number of beads is used for each reaction (∼3.2x10^5^ beads / µL stock, with 1 µL per reaction).

NOTE: Store protein-coated beads at 4 °C and use within 1 week. Make sure to thoroughly resuspend each bead preparation before using in actin assembly assays.

2.6. Determine the amount of protein coupled to the beads.

2.6.1. Pass bead suspensions for each sample through a 0.2 µm centrifugal filter.
2.6.2. Collect the beads without washing and add same volume of 1x SDS-PAGE sample buffer as the original sample volume.
2.6.3. Prepare SDS-PAGE samples containing known amounts of purified GST-CBR115 or GST-VVCA as standards and run them on the same gel to generate a standard curve.
2.6.4. Run samples on a 5-20% SDS gel.
2.6.5. Transfer protein samples to PVDF membrane following standard Western blot protocols.
2.6.6. Use a 1:1,000 dilution of anti-CARMIL1 polyclonal antibody (or anti-N-WASP polyclonal antibody) to probe the membrane overnight at 4 °C.
2.6.7. Probe with peroxidase-conjugated goat anti-rabbit IgG secondary antibody at a 1:10,000 dilution.
2.6.8. Develop blots with chemiluminescence substrate per manufacturer’s instructions.
2.6.9. Digitize autoradiograms by scanning or taking a photo. Analyze the intensities of bands using the gel analysis tool of ImageJ ^26^. Generate a standard curve by using the standards of GST-CBR115 or GST-VVCA made in step 2.6.3.

## 3. Lipid-coated bead preparation

To better mimic actin growth at a cell membrane, silica beads are coated with lipid. Choose the lipid composition of the liposomes that are fused to the beads according to the requirements of the membrane-binding proteins being studied (see the NOTE after step 3.1.1). Initially, we tested lipid mixtures based on protocols published by Taunton and colleagues ^27^ and Drubin and colleagues ^28^.

### 3.1. Making liposomes

3.1.1. Mix lipid stocks together to give a final total concentration of 5 mM at the following molar ratios: 44% DOPC, 5% DGS-NTA (Ni), 50% DOPS, and 1% Cy5-PC or 1% pyrene PE.

NOTE: The fluorescently conjugated lipids are for visualization of the lipid on the bead surface. The Ni-conjugated lipid is for coupling His-tagged proteins to the bead surface.

CAUTION: Follow all proper chemical safety instructions for using chemicals and wear appropriate personal protective equipment (PPE), such as gloves, while making liposomes. Lipid stocks are in chloroform, which should be handled in a well-ventilated area or fume hood.

3.1.2. Dry the lipid mixture under a stream of argon, and then under vacuum in a desiccator for 1 h.

3.1.3. Resuspend the lipids in MOPS buffer (step 1.2) and perform three cycles of vortexing (30 s) and bath sonication (5 min).

3.1.4. Freeze and thaw the liposomes 5 times using a dry ice/ethanol bath or liquid nitrogen.

3.1.5. Using a mini-extruder, pass the liposomes through a 0.2 µm polycarbonate membrane 9 times, and collect the extruded liposomes.

NOTE: Store prepared liposomes at 4 °C and use within 1 week. If you freeze liposomes, they must be re-extruded after thawing.

### 3.2. Coating silica beads with lipid

3.2.1. Vortex silica beads with a diameter of 2.5 µm for 30 s to resuspend beads.

3.2.1.1. Sonicate bead suspension in a water bath for 10 min.
3.2.1.2. Centrifuge for 3 min at 2,400 x g.
3.2.1.3. Wash once with MOPS buffer.

3.2.2. Coat washed beads with poly-L-lysine by incubating them in a 0.01% solution for 1 h at room temperature (RT).

3.2.2.1. Wash twice with MOPS buffer.

3.2.3. Coat the poly-L-lysine-treated beads with lipid:

3.2.3.1. Mix 3.5 µL of beads (8.0x10^6^ beads/µL, 5.6x10^5^ beads/µL final concentration) with 5 µL of 5 mM liposomes (step 3.1) in 45 µL of MOPS buffer.
3.2.3.2. Sonicate three times for 10 s and incubate for 1 h at RT with frequent gentle agitation.
3.2.3.3. Wash lipid-coated beads twice in MOPS buffer.

### 3.3. Coupling His-tagged proteins to the Ni-lipid beads

3.3.1. Incubate beads with 2 µM 6xHis-VVCA and 2 µM 6xHis peptide for 2 h on ice in MOPS buffer with frequent agitation to maintain the beads in suspension.

3.3.2. Centrifuge and wash beads once in assay buffer (no ATP) containing 1% BSA, then again in assay buffer (no ATP) containing 0.1% BSA, and finally resuspend in assay buffer containing 1 mM ATP and 0.1% BSA.

NOTE: The 6xHis peptide is used as a placeholder when not utilizing another His-tagged protein (such as His-CBR126), so that the total concentration of 6xHis is 4 µM in all samples during the coating process. Store protein-coated beads at 4 °C and use within 1 week. Make sure to thoroughly resuspend each bead preparation before using in actin assembly assays.

## 4. Actin Biomimetic Growth Assays

### 4.1. Preparation of coverslips and flow chambers

4.1.1. Acid-wash coverslips in 1 M HCl, rinse them thoroughly with water, and store them in ethanol.

4.1.2. To use a coverslip, remove excess ethanol and pass the coverslip through a flame to burn off the remaining ethanol.

4.1.3. Place two pieces of double-sided tape on a microscope slide, place the clean coverslip on top of the tape, and press gently to seal the coverslip to the tape. This creates a channel through which the sample is introduced.

4.1.4. Add the bead reaction mixture to one side of the channel until the channel is full, and then seal both ends with clear nail polish.

CAUTION: Wear appropriate PPE when acid-washing coverslips and dispose of HCl according to institutional chemical safety procedures. Take care when using an open flame to burn ethanol off the coverslips and keep the ethanol stock away from the flame.

### 4.2. Actin asymmetric growth control sample (“tails”), 20 µL

4.2.1. Prepare the following premixes separately, and incubate them for 15 min at RT so that the relevant complexes form before the actin polymerization reaction starts:

4.2.1.1. 0.9 µL of 22.73 µM Arp2/3 complex (100 nM final concentration) with 1 µL of VVCA-coated beads (∼3.2x10^5^ beads per reaction; ∼1.6x10^4^ beads/µL final) and CP (50 nM final concentration) in assay buffer (+ATP) containing 2 µL of 10x β-ME and 2 µL of 10x BSA + methyl cellulose.
4.2.1.2. Actin monomers (5 µM final concentration; a mixture of unlabeled and Alexa 488-labeled actin, see step 4.2.2) with profilin (5 µM final concentration).

4.2.2. Calculate the volumes of unlabeled and labeled actin so that 10% of the final amount of actin is Alexa 488-labeled, as follows:

4.2.2.1. Calculate the total nanomoles of actin for 5 µM in 20 µL (0.1 nmoles).
4.2.2.2. For labeled actin, multiply the total nanomoles of actin by the desired labeled fraction (0.1).
4.2.2.3. Divide that value by the fraction of labeling of the Alexa-488 actin (e.g., if the percentage of labeling is 50%, divide by 0.5) to get nanomoles of labeled actin.
4.2.2.4. Multiply the nanomoles of labeled actin by 1000, then divide by the stock concentration (in µM) to obtain the volume (in µL) to add to the mixture.
4.2.2.5. For nanomoles of unlabeled actin, take the total nanomoles and subtract the value of the nanomoles of labeled actin.
4.2.2.6. Multiply the nanomoles of unlabeled actin by 1000, then divide by the stock concentration (in µM) to obtain the volume (in µL) to add to the mixture.

4.2.3. Combine all the premixes in one 1.5 mL microcentrifuge tube (20 µL final volume) to start the polymerization reaction, and incubate for 15 min at RT.

4.2.4. Load the reaction mixture into a flow chamber, seal it with clear nail polish (step 4.1.4), and image after another 15 min.

NOTE: Beads will have asymmetric actin growth (“tails”) after 30 min of reaction time. These reaction times were chosen after pilot experiments testing a range of times.

### 4.3. Actin growth in the presence of CP regulators, 20 µL

4.3.1. Incubate His-VVCA beads with various concentrations of untagged CBR126 (a longer portion of CARMIL1 that includes the full membrane binding domain) for 15 min at RT.

4.3.2. Centrifuge and resuspend in assay buffer with 0.1% BSA and 1 mM ATP.

NOTE: We confirmed that CBR126 remained associated with lipid beads over the course of several hours using fluorescently labeled CBR126, but we always used the beads within a few days for the assays reported here. One can also use GST-CBR115 and glutathione-functionalized beads (section 2) in place of the Ni-lipid beads (section 3).

4.3.3. Prepare the following premixes separately, and incubate them for 15 min at RT so that the relevant complexes form before the actin polymerization reaction starts:

4.3.3.1. 0.9 µL of 22.73 µM Arp2/3 complex (100 nM final concentration) with 1 µL of VVCA- and CBR126-coated beads (∼3.2x10^5^ beads per reaction; ∼1.6x10^4^ beads/µL final) in assay buffer (+ATP) containing 2 µL of 10x β-ME and 2 µL of 10x BSA + methyl cellulose
4.3.3.2. Actin monomers (5 µM final concentration; a mixture of unlabeled and Alexa 488-labeled actin, see step 4.2.2) with profilin (5 µM final concentration).
4.3.3.3. CP (50 nM final concentration) with V-1 (500 nM final concentration).

4.3.4. Combine all the premixes in one 1.5 mL microcentrifuge tube (20 µL final volume) to start the polymerization reaction, and incubate for 15 min at RT.

4.3.5. Load the reaction mixture into a flow chamber (step 4.1), seal it with clear nail polish, and image after another 15 min.

NOTE: Alternatively, arrest the reaction after 30 min by adding an equal volume of 2x arrest mixture (step 1.3), load the sample into a flow chamber, and image immediately.

### 4.4. Two-color actin bead assays

4.4.1. Make bead protein mixtures including 50 nM CP, 100 nM Arp2/3 complex, 5 µM profilin, 5 µM Alexa488-actin and 500 nM V-1 (where indicated) as described in step 4.3.

4.4.2. Allow reactions to proceed for 5 min.

4.4.3. Dilute the reaction 1:5 into an identical protein mixture containing 5 µM Alexa594-actin in place of Alexa488-actin.

4.4.4. After 5 min (or longer if needed) arrest the reaction by adding an equal volume of 2x arrest mixture (step 1.3).

4.4.5. Load the sample into a flow chamber (step 4.1), and image immediately.

### 4.5. Antibody labeling of beads

Antibodies can be used to determine the localization and/or amount of protein attached to the bead or within the actin network. Perform these steps after the bead assays in sections 4.2-4.3.

4.5.1. Prepare bead samples as described in sections 4.2-4.3.

4.5.2. At 30 min, arrest the actin polymerization reactions by adding an equal volume of 2x arrest mixture (step 1.3).

4.5.3. Pellet the beads by centrifugation (2,400 x g, 3-5 min) and resuspend them in assay buffer (no ATP) containing 10 µM phalloidin, 3% BSA, and anti-CARMIL1 or anti-Arp3 primary antibody (1:400). Incubate for 2 h at RT with frequent, gentle agitation or on a rotator.

4.5.4. Wash twice using assay buffer (no ATP) containing 10 µM phalloidin.

4.5.5. Add Alexa 647-labeled goat anti-rabbit IgG or Alexa 647-labeled donkey anti-mouse IgG secondary antibody (1:400) in assay buffer (no ATP) containing 10 µM phalloidin and 3% BSA. Incubate for 1 h at RT with frequent, gentle agitation or a rotating block.

4.5.6. Wash beads twice using assay buffer (no ATP) containing 10 µM phalloidin.

4.5.7. Resuspend in assay buffer (no ATP) + 20 mM βME + 2.5 mg/mL BSA + 0.2 % methyl cellulose + 10 µM phalloidin, and load sample into coverslip chamber.

4.5.8. Image samples as described in section 5.

### 4.6. Alternative NPF constructs as activators of Arp2/3 complex

Actin bead assays utilize portions of the VCA domain of N-WASP or a similar NPFs to activate Arp2/3 complex. Using a longer fragment, termed here PVVCA, includes a proline-rich domain that binds profilin.

4.6.1. Incubate beads with 2 µM 6xHis-PVVCA and 2 µM 6xHis peptide for 2 h on ice in MOPS buffer with frequent agitation to maintain the beads in suspension.

4.6.2. Centrifuge and wash beads once in assay buffer (no ATP) containing 1% BSA, then again in assay buffer (no ATP) containing 0.1% BSA, and finally resuspend in assay buffer containing 1 mM ATP and 0.1% BSA.

4.6.3. Prepare the following premixes separately, and incubate them for 15 min at RT so that the relevant complexes form before the actin polymerization reaction starts:

4.6.3.1. 0.9 µL of 22.73 µM Arp2/3 complex (100 nM final concentration) with 1 µL of PVVCA-coated beads (∼3.2x10^5^ beads per reaction; ∼1.6x10^4^ beads/µL final) and CP (50 nM final concentration) in assay buffer (+ATP) containing 2 µL of 10x β-ME and 2 µL of 10x BSA + methyl cellulose
4.6.3.2. Actin monomers (5 µM final concentration; a mixture of unlabeled and Alexa 488-labeled actin, see step 4.2.2) with profilin (5 µM final concentration).

4.6.4. Combine all the premixes in one 1.5 mL microcentrifuge tube (20 µL final volume) to start the polymerization reaction, and incubate for 15 min at RT.

4.6.5. Load the reaction mixture into a flow chamber (step 4.1), seal it with clear nail polish, and image after another 15 min.

## 5. Microscope imaging

Once bead samples are loaded into coverslip chambers, they can be imaged at RT using wide-field fluorescence microscopy. NOTE: Allow the nail polish to dry for a few minutes before placing the slide on the microscope.

5.1. Use a 60x 1.45 NA oil objective on an inverted wide-field fluorescence microscope with a mercury lamp for fluorescence excitation.

5.2. Image fluorescent actin using a GFP (for Alexa-488) or a Texas-Red (for Alexa-594) filter set. Use the Texas-Red filter set for fluorescently labeled CP or CBR126. Use the Cy5 or DAPI/pyrene filter set for imaging fluorescent lipid on the bead surface. Collect a bright-field image as well (to observe the bead).

NOTE: When imaging multiple fluorescent channels, confirm that there is no bleed-through from one channel into another. To test this, image samples containing each fluorescent component individually in every channel. Image the far-red channel first and proceed toward the shortest-wavelength channel.

5.3. Collect 16-bit images with an EM-CCD camera using Micro-Manager software^29^.

5.4. Process digital images using Fiji software ^30^.

## 6. Analysis of fluorescence area, circularity, and intensity around beads

Use Fiji, an ImageJ analysis platform, to process all digital images.

### 6.1. Actin fluorescence area

In actin fluorescence images, there are two areas of fluorescence: the area of brightest fluorescence near the bead surface and the total area including the diffuse cloud of actin.

6.1.1. For the area of brightest fluorescence near the bead surface: outline the area using the freehand drawing tool and the Measure tool in Fiji ^30^.

6.1.2. For the area when beads are surrounded by diffuse clouds of actin fluorescence: set a threshold level based on the fluorescent signal under Image>Adjust>Threshold. Use the mask created by the threshold to demarcate the edge of the actin cloud. Use the freehand drawing tool to outline around the threshold mask to determine the area of the actin cloud.

### 6.2. Actin fluorescence shape

6.2.1. Circularity: determine the circularity of the fluorescence for the area of brightest fluorescence closest to the beads.

6.2.1.1. Use the Freehand tool to draw around the area of brightest fluorescence.
6.2.1.2. Use the Measure tool to determine the circularity of each shape.

6.2.2. Roundness: determine the roundness of the fluorescence for the area of brightest fluorescence closest to the beads.

6.2.2.1. Use the threshold tool to determine the area of brightest fluorescence closest to the bead surface and set a threshold mask.
6.2.2.2. Use the Freehand tool to draw around the area of the threshold mask.
6.2.2.3. Use the Measure tool to determine the aspect ratio and roundness of each shape.

NOTE: Circularity is a metric within the Measure tool (Analyze>Measure) defined by ImageJ as circularity = 4pi(area/perimeter^2), with the value of 1 indicating a perfect circle. Roundness is a metric within the Measure tool (Analyze>Measure) defined by ImageJ as the inverse of the aspect ratio, with the value of 1 indicating a perfect circle. To include Circularity or Roundness measurements, select “Shape Descriptors” under Analyze > Set Measurements.

### 6.3. Total actin, CP, or CBR fluorescence

6.3.1. Draw an ∼40x40 µm box centered around a bead.
6.3.2. Use the Measure tool to obtain the value of the integrated density.
6.3.3. Draw an ∼50x50 µm box around the outside of an ∼40x40 µm box to determine the background value surrounding each bead.

6.3.3.1. Calculate the background using the following equation:

F_background_ = (F_o_ - F_i_)*(1615 µm^2^/(2512 µm^2^ - 1615 µm^2^)) Note: 1615 and 2512 correspond to the areas of the smaller and larger boxes, respectively.

NOTE: In this equation, F_o_ is the outer box fluorescence, and F_i_ is the inner box fluorescence.
6.3.4. Calculate the total fluorescence of the bead as:

F_tot_ = F_i_ – F_background_

### 6.4. Radial variance, radial intensity, and variance metric

NOTE: For these image analysis parameters, we use custom code that we have made freely available at https://zenodo.org/records/13765359. The code is written in Python and uses the PIL, NumPy, and scikit-image libraries. The details for the methods and formulas used in the original code can be found in reference ^19^.

6.4.1. Manually crop TIFF images of beads to isolate beads in focus.
6.4.2. Use the code deposited above to analyze the TIFF images.

## Results

Capping protein (CP) is necessary for Arp2/3-based actin to grow asymmetrically away from the bead, resulting in “tails” of filamentous actin ^12, 31^. Using glutathione-functionalized beads containing GST-VVCA, we observed a biphasic response, where too little CP resulted in little actin growth, and actin growth was also inhibited at high CP concentrations. The most robust actin growth using our bead assay protocol was observed with 50 nM CP and 100 nM Arp2/3 complex (Figure 1, panel A). The complete absence of CP resulted in bright actin fluorescence immediately surrounding the bead along with a diffuse actin “cloud” farther from the bead surface. The area of fluorescence and total actin fluorescence were quantitated as described in section 6 (Figure 1, panels B and C).

**Figure 1.**
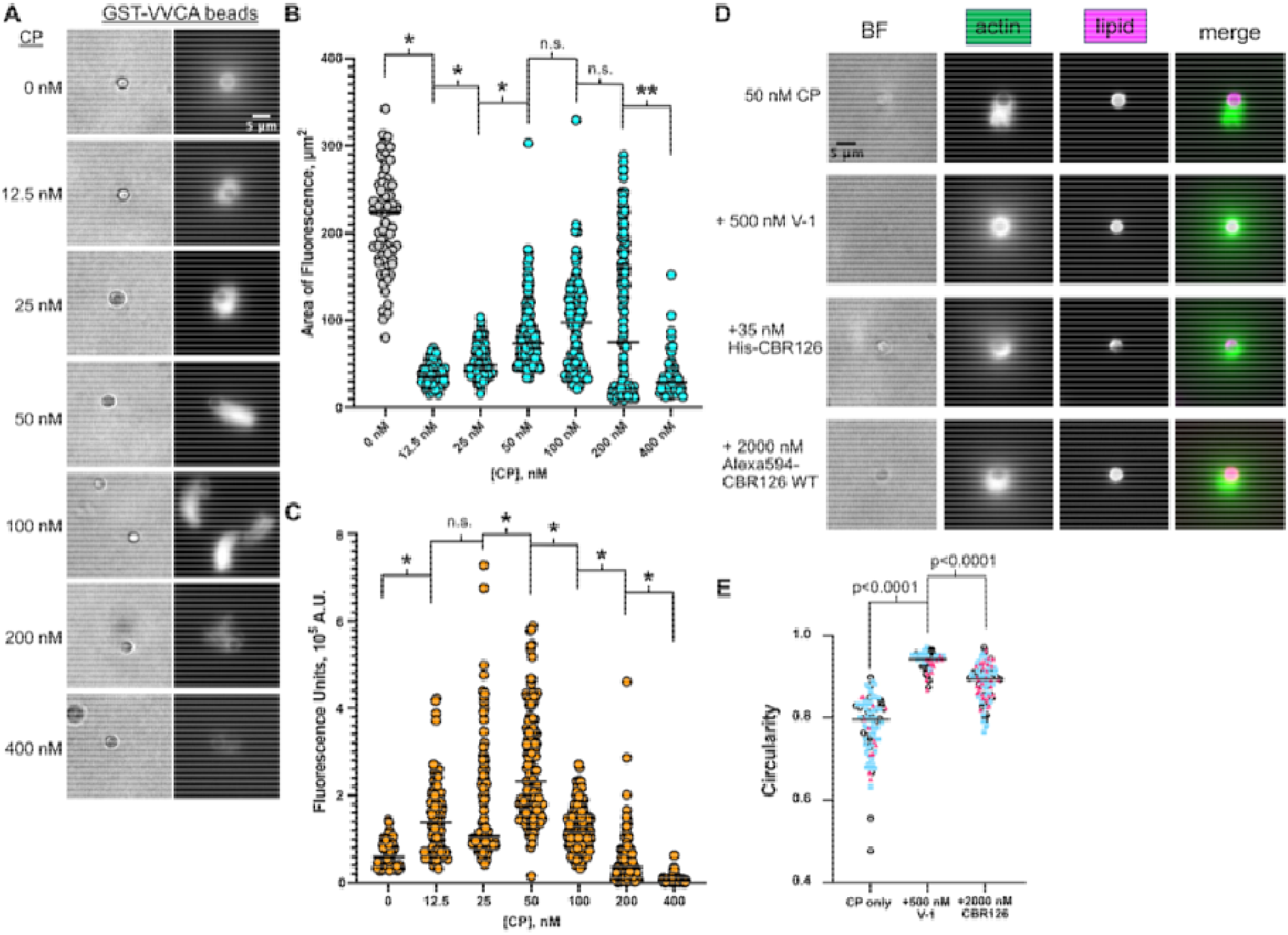
Examples of bead assay results using glutathione-functionalized beads or lipid-coated Ni-functionalized beads. **A**. Actin growth assays on glutathione-functionalized beads containing GST-VVCA, with mixtures of 100 nM Arp2/3 complex, 5 µM profilin-actin, and varied CP concentrations, as shown. Brightfield (BF) is shown on the left and actin fluorescence (Alexa488-actin) is the right column. Images were collected 30 min after mixing. Scale bar 5 µm. **B**. The area of fluorescence was collected and plotted with each dot representing a bead. The grey dots represent the total area of the fluorescence, including the actin “cloud.” The cyan dots represent the area of the asymmetric actin growth, and those samples did not contain “clouds.” Data from four independent experiments on different days are shown. The horizontal black line shows the median. P values were calculated from non-parametric Mann-Whiteney t-tests, where * is p<0.0001, ** is p>0.01, and n.s. is not significant. **C**. Total actin fluorescence was calculated as described in section 6. Each dot represents one bead. Data from four independent experiments on different days are shown. The horizontal black line shows the median. P values were calculated from non-parametric Mann-Whiteney t-tests, where * is p<0.0001 and n.s. is not significant. **D**. Actin network growth on Ni-functionalized lipid beads containing His-VVCA was similar to using glutathione beads. Images of beads were collected after 30 min. Lipids are visualized using Cy5-PC. Scale bar 5 µm. **E**. Circularity is plotted for each bead. Three independent experiments are plotted as different colors and shapes. The horizontal black line shows the median. P values shown were calculated from non-parametric Mann-Whiteney t-tests. Panels A-C are adapted from Mooren et al. (6). Panels D-E are adapted from Mooren et al. (5).

We observed similar results to glutathione-functionalized beads when we applied Ni-lipid to the bead surface and allowed actin filament networks to assemble utilizing the same protein components (Figure 1, panel D, top row). Adding stoichiometric amounts of V-1 to the mixture completely inhibited CP activity, and beads resembled samples without CP (Figure 1, panel D, second row). When a fragment of CARMIL1, containing the region necessary and sufficient for CP binding and activation, termed CBR126, was attached to the bead surface via the Ni-His interaction, a small amount of asymmetric actin growth was observed from the bead surface ^18^ (Figure 1, panel D, third row). However, when CBR126 was targeted to the bead surface via its membrane-binding domain, more robust asymmetric actin growth was observed (Figure 1, panel D, last row). Circularity was a quantitative metric for assessing the level of asymmetric actin growth, as shown in Figure 1, panel E.

To attach lipid to our silica beads we first tested various lipid ratios based on published methods ^27, 28^ of either 94:0:5:1 or 74:20:5:1 DOPC:DOPS:Ni-DGS:Cy5PC in our actin assembly assays. We attached His-VVCA to the beads, and found that, with 100 nM Arp2/3 complex, 50 nM CP, and 5 µM profilin-actin, asymmetric actin growth was most robust on 74:20:5:1 lipid-coated beads (Figure 2, panel A). However, by performing liposome sedimentation assays with our membrane-binding protein fragment, CBR126, we determined that a higher ratio of DOPS (50%) was necessary for effective interaction of the protein with the lipid (see methods section of reference ^18^ for the protocol for sedimentation assays). Utilizing the fluorescence of the Cy5 PC, we observed that the higher ratio DOPS lipid (44:50:5:1 DOPC:DOPS:Ni-DGS:Cy5PC) did not coat the beads well (Figure 2, panel B). Therefore, we attempted various methods to make the silica beads “sticky” for the higher PS ratio liposomes using various amounts of (3-aminopropyl)triethoxysilane (APTES) on the silica beads by incubating the beads in APTES solutions for 1 h at RT. This treatment made the liposomes adhere well to the beads but did not give uniform coverage of the bead surface, as demonstrated by the patchy fluorescence observed (Figure 2, panel C, rows 1-4) and tubules that were often emanating from the bead surface. Next, we treated the silica beads with poly-L-lysine by incubation for 1 h at RT. The beads coated with poly-L-lysine displayed uniform fluorescence, indicating a more uniform coverage of the lipid onto the bead surface (Figure 2, panel C, bottom row). We confirmed that poly-L-lysine treatment of the beads yielded good coverage of the lipid on the bead surface by scanning electron microscopy (Figure 2, panel D).

**Figure 2:**
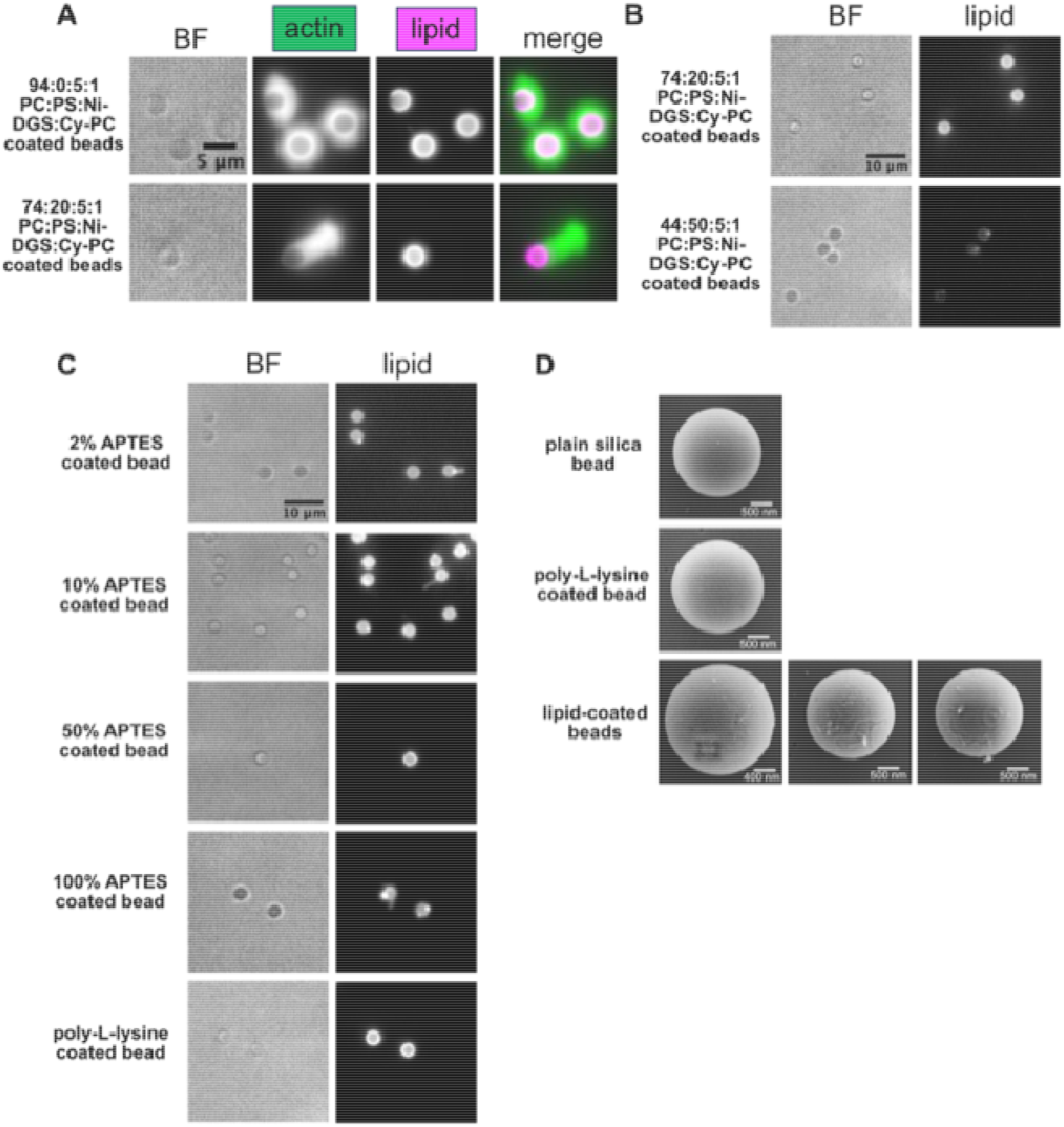
Optimization of Ni-functionalized lipid coating of silica beads. **A**. Beads were coated with lipids comprised of either 94:0:5:1 or 74:20:5:1 DOPC:DOPS:Ni-DGS:Cy5PC in our actin assembly assays. BF, Alexa488-actin, and Cy5-lipid fluorescence are shown. Scale bar 5 µm. **B**. Beads were coated with either 74:20:5:1 or 44:50:5:1 DOPC:DOPS:Ni-DGS:Cy5PC and the Cy5-lipid fluorescence was collected. Scale bar 10 µm. **C**. Increasing amounts of APTES was used to treat the silica beads prior to incubation with 44:50:5:1 DOPC:DOPS:Ni-DGS:Cy5PC liposomes (rows 1-4). Poly-L-lysine was used to treat the silica beads prior to incubation with 44:50:5:1 DOPC:DOPS:Ni-DGS:Cy5PC liposomes (bottom row). Scale bar 10 µm. **D**. Scanning electron micrographs of silica beads that were washed (top row), coated with poly-L-lysine (middle row), and coated with 44:50:5:1 DOPC:DOPS:Ni-DGS:Cy5PC liposomes (bottom row, three examples of different beads). Taken from reference (Mooren et al JBC 2026). Scale bars as indicated, 400 and 500 nm.

Arp2/3 complex is inactive on its own and must be activated by an NPF. Activation of Arp2/3 complex by WASP-family proteins is accomplished through a region termed the VCA domain, encompassing the WASP-homology 2 (WH2)/Verprolin-homology (V), central (C), and acidic (A) domains ^22^ Two V domains in tandem provide more activation than does one ^32–34^. Verprolin homology domain is also known as WH2 for Wasp Homology 2 ^22, 23^. Structural studies demonstrate that these regions induce a conformational change in Arp2/3 complex, allowing optimal binding to a mother filament and nucleation of a new actin branch with a free barbed end that will continue to grow. NPFs also recruit new actin monomers to Arp2/3 complex. This is accomplished, in part, by binding to profilin-actin through a proline-rich domain (PRD) upstream of the VCA domain ^35^.

N-WASP is an NPF that contains a PRD, followed directly by two tandem WH2 domains; including the PRD enhances Arp2/3 branching / nucleating activity ^7, 35^. Typically, in our assays we have used an N-WASP fragment that consists solely of the two WH2 domains, without the N-terminal PRD. To examine the effect of the PRD upstream of the two WH2 domains in our assays, we generated and purified a His-tagged PVVCA fragment, attached it to Ni-lipid coated beads and compared the resulting actin growth to results with His-VVCA. Some beads with His-PVVCA grew very long tails in the same amount of time compared to His-VVCA beads (Figure 3). This result confirms previous ones^35^. The presence of the PRD can then be used to fine-tune the strength of Arp2/3 activation and actin assembly. For the purposes of our experiments investigating CP regulators, we found that having less robust actin assembly, using the His-VVCA, facilitated observations of differences in the actin network organization.

**Figure 3.**
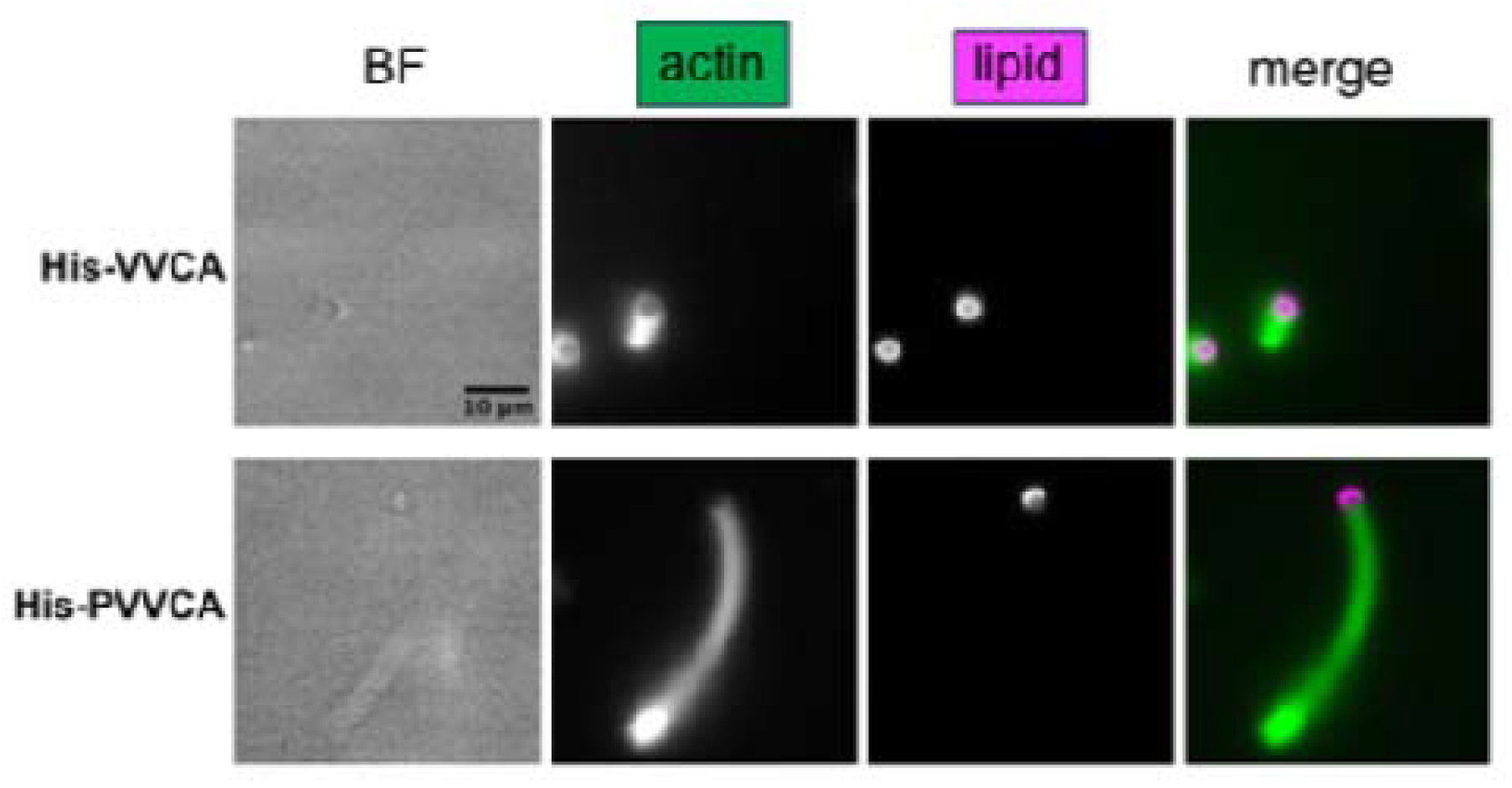
Testing NPF domains in actin bead assays. Actin networks were generated for 30 min on either His-VVCA-or His-PVVCA-coated Ni-lipid beads using 100 nM Arp2/3 complex, 50 nM CP, and 5 µM profilin-actin. BF, Alexa488-actin, and Cy5-PC fluorescence is shown. Scale bar is 10 µm.

To visualize where actin barbed ends were oriented and where the actin growth was occurring in our bead samples, for example, at the bead surface, we utilized two different fluorescently labeled actin mixtures, adding them at different times. Alexa488-actin was included in the initial reaction and then replaced by Alexa594-actin after 5 min of reaction time. Samples were then arrested using a mixture of latrunculin and phalloidin to stabilize the actin and prevent further growth. The results showed that the new actin growth was occurring at the bead surface (Figure 4, panel A), as has been previously reported ^19, 36^.

Visualizing the localization of proteins of interest within the actin network generated in bead assays was possible by using fluorescently conjugated proteins or by using specific antibodies. For example, we observed the localization of Arp2/3 complex in or actin bead assays by using an anti-Arp3 antibody and a fluorescently tagged secondary antibody (Figure 4, panel B).

**Figure 4.**
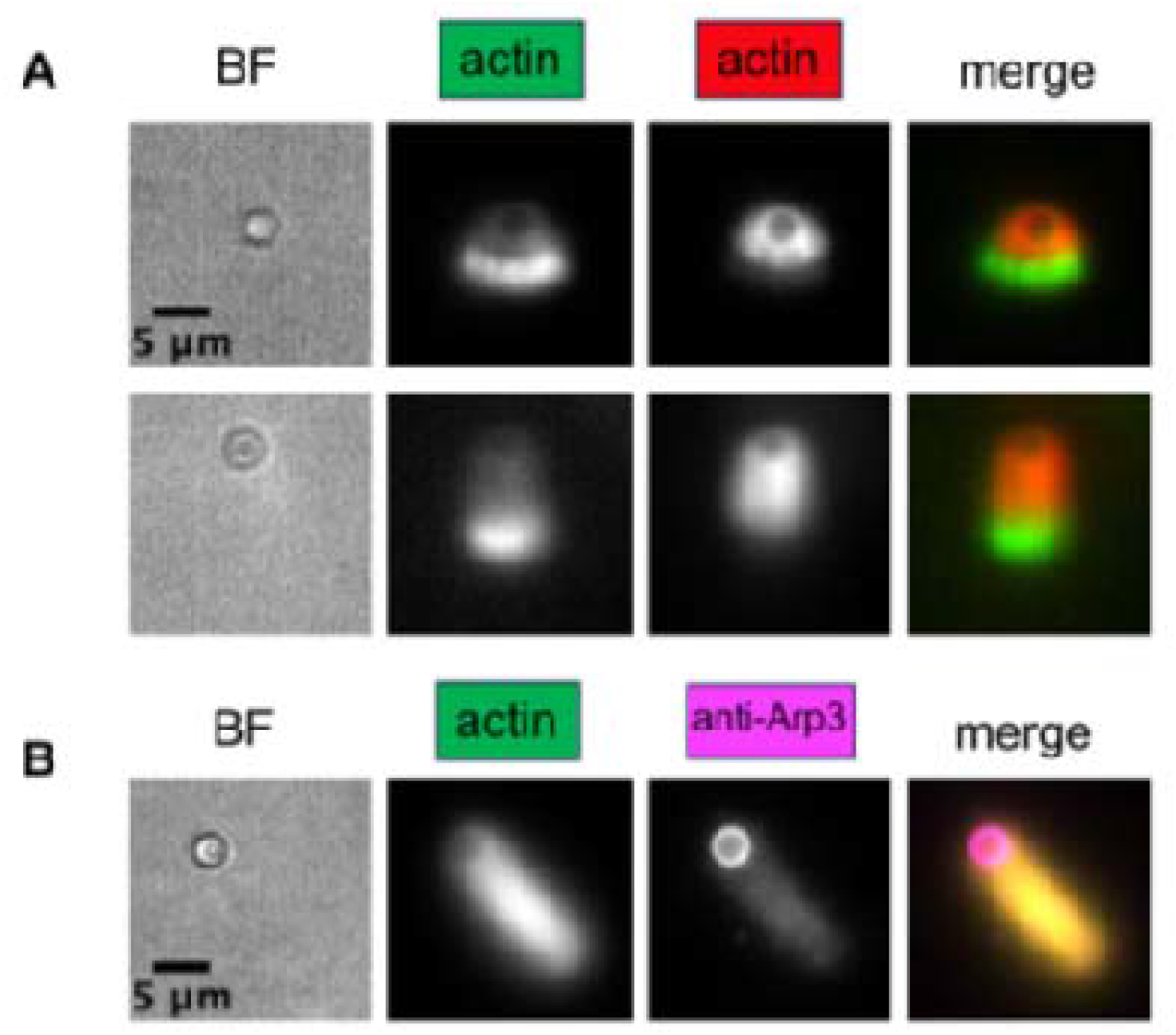
Localization of actin and Arp2/3 complex around beads. Scale bars are 5 µm. A. Two-color actin experiment. Actin networks were generated for 5 min by incubating 100 nM Arp2/3 complex, 50 nM CP, and 5 µM profilin-actin using Alexa488-actin. Then the beads were diluted into an identical protein mixture containing Alexa594-actin. Reactions were arrested after 5 min and imaged. B. Arp2/3 antibody-labeling of actin networks. Actin networks were generated by incubating 100 nM Arp2/3 complex, 50 nM CP, and 5 µM profilin-actin using Alexa488-actin. Reactions were arrested after 30 min. Anti-Arp3 antibody was incubated with arrested bead samples, followed by incubation with Alexa647 secondary antibody and imaged. Figure adapted from Mooren et al. (6).

## Discussion

Reconstitution of actin assembly is a well-established approach for dissecting the molecular mechanisms of actin-based motility ^12, 15, 36, 37^. Historically, actin filament assembly and actin-based motility were first observed in association with bacteria in cytoplasm of living cells and in whole-cell extracts ^38, 39^. In those settings, actin assembly required Arp2/3 complex and an NPF ^40^. Carlier and colleagues made the landmark discovery that actin assembly and motility could occur on the bacteria placed into synthetic biochemical reconstitutions systems composed of purified proteins, including Arp2/3 complex, ADF/cofilin, and CP ^12^. Later, their group demonstrated that assembly and movement could be reconstituted with carboxylated polystyrene beads in the synthetic mixture ^37^.

These studies and many later ones have used passive adsorption of proteins to the bead surface ^15, 37^. For our experiments investigating regulation of CP, we needed to target the CP activator CARMIL and an NPF to the bead surface to mimic their localization to the cell membrane in cells. To this end, we developed two methods for protein attachment to the bead surface. The first method utilizes glutathione-functionalized polystyrene beads and GST-fusion proteins. This method allows for consistent control over the amounts of each protein by keeping the total molar amount of GST constant, including GST-fusion proteins and plain GST to saturate the glutathione residues of the bead. By normalizing the number of beads added to each reaction by OD600, this approach allows for consistent formation of actin networks because the amounts of NPF and CARMIL available for activation of Arp2/3 complex and CP, respectively, are relatively constant among individual beads. Implementing these procedures allows for careful fine-tuning of proteins to study their contributions to the growth and organization of an actin network.

The second method we developed is to coat silica beads with supported lipid bilayers by fusing liposomes to the beads, following the methods of Taunton and colleagues ^13^. With this approach, the relative amounts of lipids on the beads can modified as needed for individual experiments. This approach couples proteins to the lipid on the bead surface with nickel coupled lipids and His-tagged proteins ^27^ and the total molar amounts of 6xHis is held constant as in the GST/glutathione bead experiments. Or a membrane-binding protein of interest can interact directly with the lipids on the bead. Most important, the proteins can move laterally on the bead surface, as they would in a membrane ^41^.

The lipid composition must suit the membrane-binding proteins being studied, and the bilayer must cover the bead uniformly. We initially used a ratio of 74:20:5:1 DOPC:DOPS:Ni-DGS:Cy5PC (Figure 2A) to coat the beads. This lipid composition worked well in the absence of CP regulators and when we added CARMIL via the His-tag/Ni-lipid interaction. We also tested the interaction of the intrinsic membrane-binding domain of the CP-activating fragment of CARMIL, which is found in tandem with the CP-binding CPI and CSI motifs with liposomes and needed to use a higher percentage of PS for efficient binding ^18^. Therefore, we made liposomes using a ratio of 44:50:5:1 DOPC:DOPS:Ni-DGS:Cy5PC. Initially, lipids with this higher content of PS did not coat the silica beads well (Figure 2B). However, we tested ways to modify the surface of the beads and to promote lipid attachment, and we found that poly-L-lysine coating provided a uniform coverage of the 44:50:5:1 DOPC:DOPS:Ni-DGS:Cy5PC lipid mixture on the beads (Figure 2C and D). Additionally, the percentage of Ni-conjugated lipid can be adjusted if needed. For our experiments, we determined that 5% of the Ni-conjugated lipid was sufficient; higher levels would allow for a greater density of surface-bound proteins. The composition of lipids can also be controlled. We used a basic mixture of DOPC and DOPS, which does not reflect the complex composition of lipids in cell membranes. A more physiological mixture of lipids (including components such as PE, PI, and cholesterol) can be used to coat silica beads. In addition, the saturation status of the lipid can also be controlled, with PO-lipids replacing DO-lipids. The protocols presented here should provide a starting point for optimization, and using a fluorescently labeled lipid will help in verifying uniform coverage of silica beads so that the composition of the mixture and the ratios of lipids can be adjusted for different experiments.

Choosing which regions of an NPF to use in bead assays deserves consideration. Recent discoveries have demonstrated how NPFs alter Arp2/3 complex conformation via the VCA domain to activate it for binding to a mother filament to nucleate a new branch (refs). Additionally, a PRD is present upstream of one or more WH2-domains that binds to profilin-actin to promote branched actin assembly ^35^. Including the PRD on our N-WASP fragment dramatically increased the amount of actin growth from the bead surface (Figure 3). This may be useful for certain applications. For our purposes when studying the role of CP regulation, having such robust actin assembly obscured the results we observed in actin network organization when we used the N-WASP fragment lacking the PRD. However, optimizing the system using the fragment that contains the PRD is of interest and should be considered in setting up bead assays in the future.

Localization of a protein of interest is also important for understanding its role in actin assembly. Utilizing fluorescent tags, such as by coupling fluorophores covalently through reactions with amino acids within the protein (i.e. maleimide reactions with a cysteine) or by creating a fluorescent protein-fusion, can be very useful in localization studies. Additionally, arresting the actin network and using antibodies to probe for the protein of interest is possible (Figure 4B) and is outlined in section 4.5. We present a method for performing two-color actin experiments in section 4.4 (Figure 4A). This method allows for one to observe where actin assembly is taking place and where barbed ends of actin filaments are located. This type of experiment can be very informative in understanding how a protein of interest contributes to actin network assembly. Observing how a protein contributes to actin network assembly and its localization within the network in a controlled subset of proteins is a powerful way to contribute to understanding its cellular function.

Several factors should be considered when interpreting the results of bead actin assembly assays. First, fluorescence intensity is a relative, uncalibrated measure of the amount of actin or of a labeled protein, so comparisons should be made only among samples imaged with identical settings and analyzed in the same way. Second, choosing which fluorophore dyes to use is important. Any combination of dyes can be coupled to proteins of interest by using chemistry between specific amino acids. For example, our CARMIL fragment, CBR126, has one cysteine located in a position that did not affect binding to CP and was ideal for using a maleimide Alexa594 to label. Careful design can allow for easy labeling of proteins using similar methods. In our experiments, Alexa488 was used for actin and Alexa594 for other labeled components, but these are FRET pairs, and additionally, Alexa594 fluorescent signal can bleed over into the far-red channel (i.e. Cy5 or Alexa647-conjugated antibodies). Imaging from far-red to blue and including singly labeled controls reduces these potential problems, and choosing a different Alexa dye, such as Alexa568, may avoid some of them; this choice will be important for experiments that require accurate quantitation of each component. Third, our measurements are made on images collected at a single time point, often after the reaction has been arrested, so they report network organization rather than assembly kinetics; time-lapse imaging, as used in comet-tail motility assays ^14^, could add kinetic information. Lastly, wide-field fluorescence microscopy does not resolve actin network ultrastructure, just course organization information. A technique such as electron microscopy would be informative about branch and filament density.

These methods also open several directions for future work. The data sets produced by these experiments are well-suited for molecular modeling studies, another area in which great progress has been made ^42–44^. Molecular modeling is a powerful method for understanding how each component contributes to the overall growth and architecture of the actin network and comparing modeling data with experimental data is necessary for interpretation of modeling. Calibrated fluorescence measurements will be especially valuable for this purpose. This is a promising area of the field that should continue to be explored. With appropriate validation, the approach can also be extended to other membrane-associated regulators, including other CPI-motif proteins ^24, 25^, and to mutant or disease-associated variants. Overall, these methods provide precise control over the identity, amount, and mode of membrane attachment of the proteins that regulate Arp2/3 complex and CP, allowing individual components to be dissected mechanistically in ways that are difficult to achieve in cells.

## Acknowledgements

We are grateful to Jennifer Obermann, James DeBrecht, and Dr. Mike Onken for advice and assistance. This work was supported by NIH grant R35 GM144082.

## Table of Materials

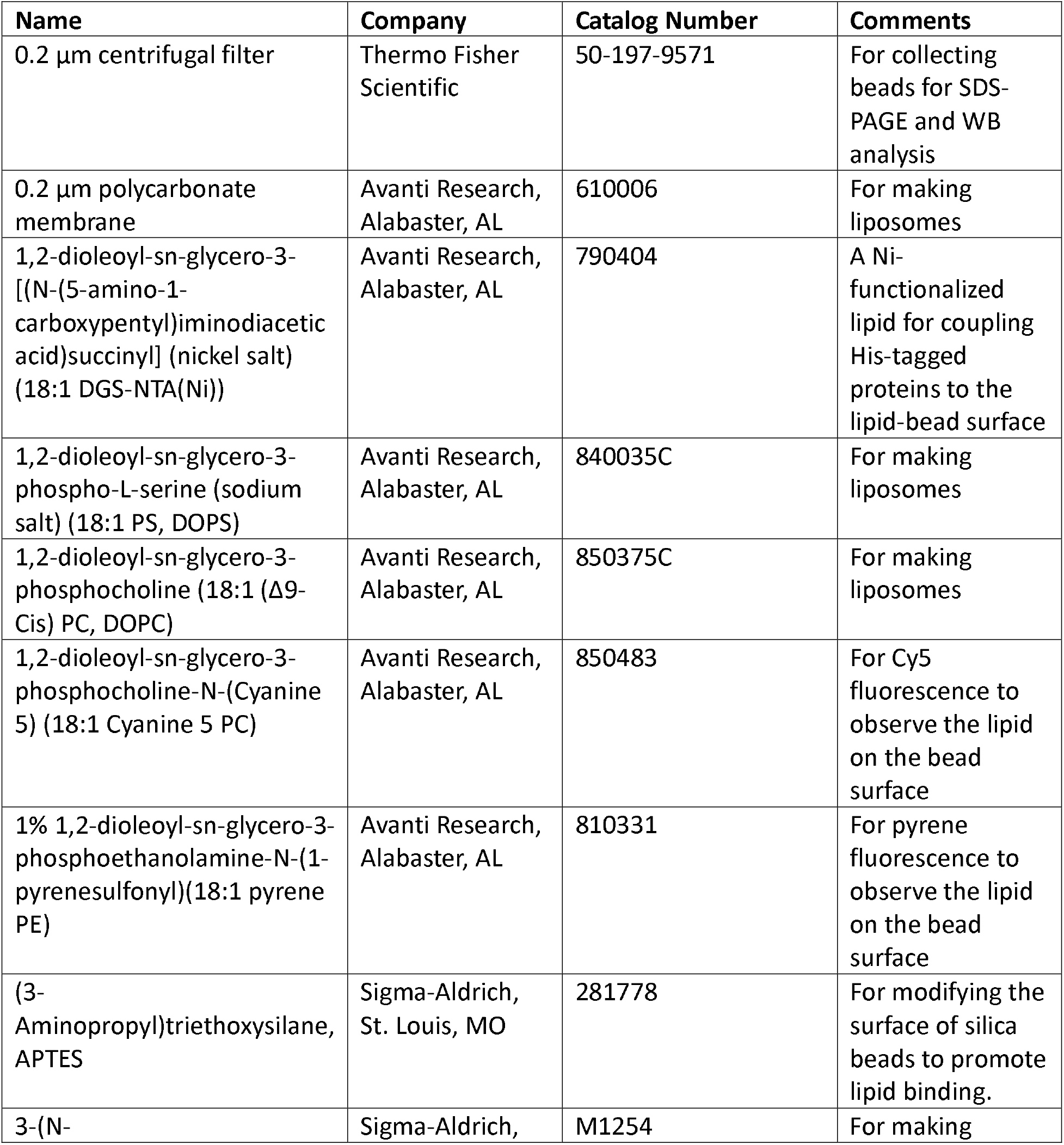
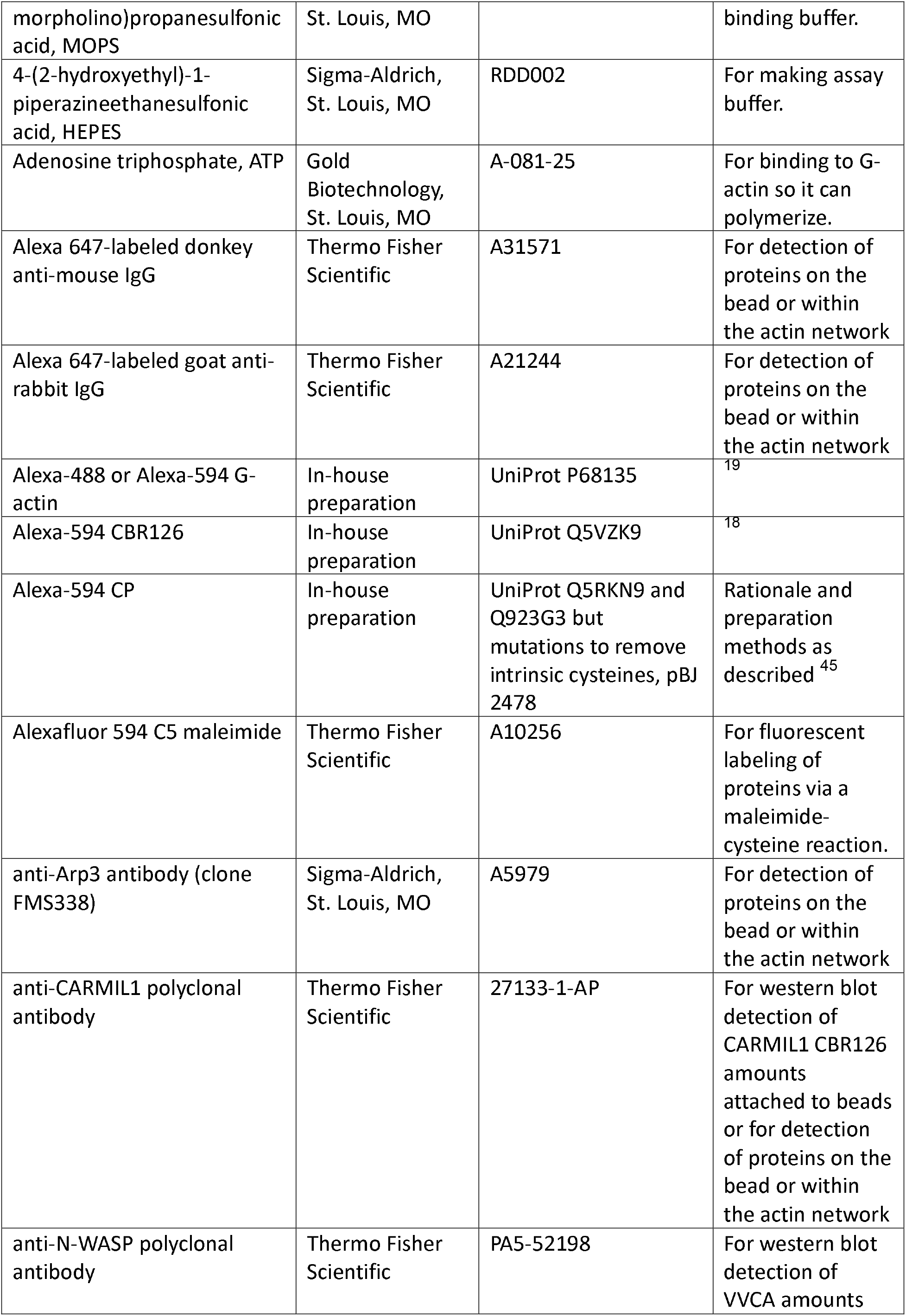
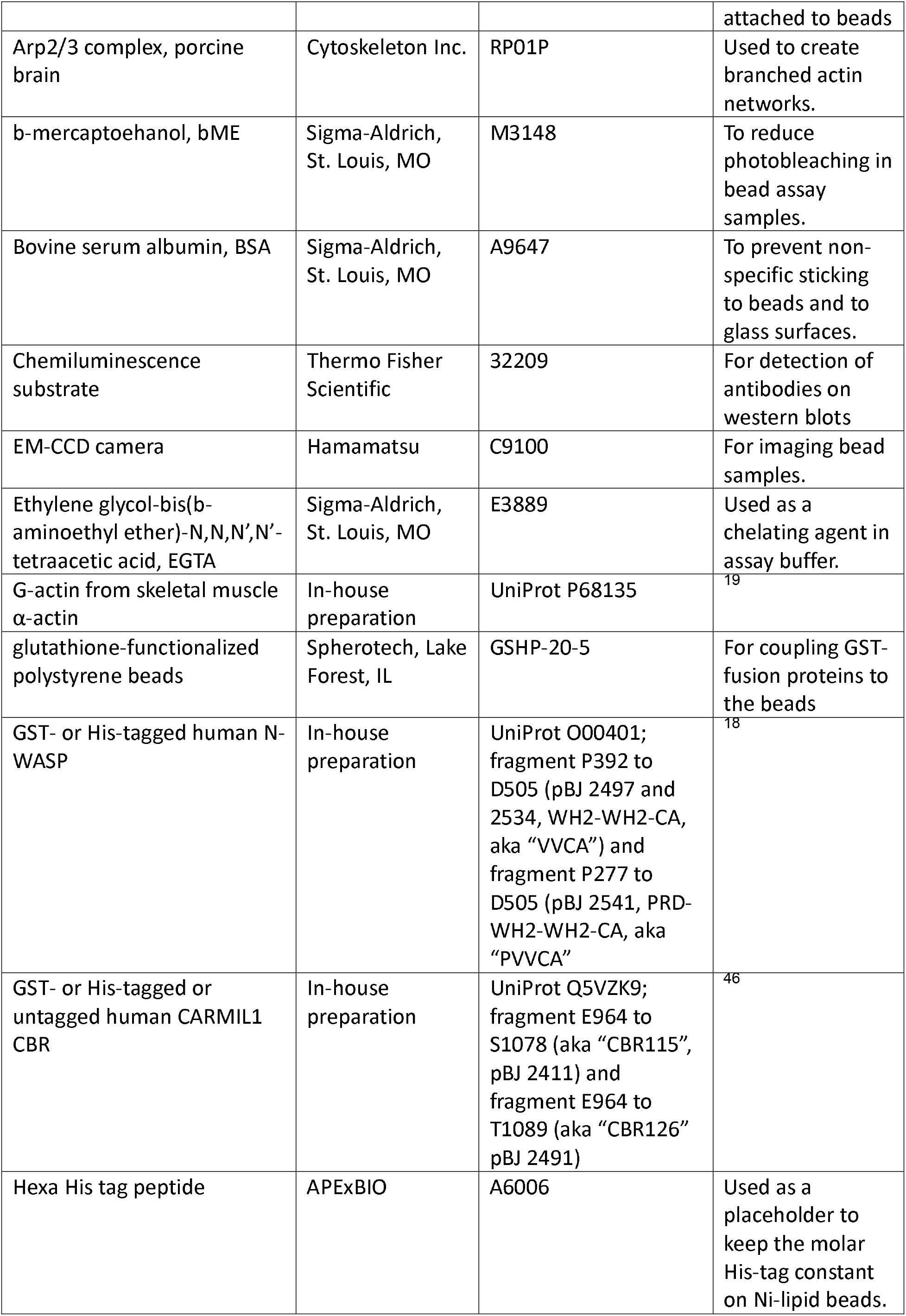
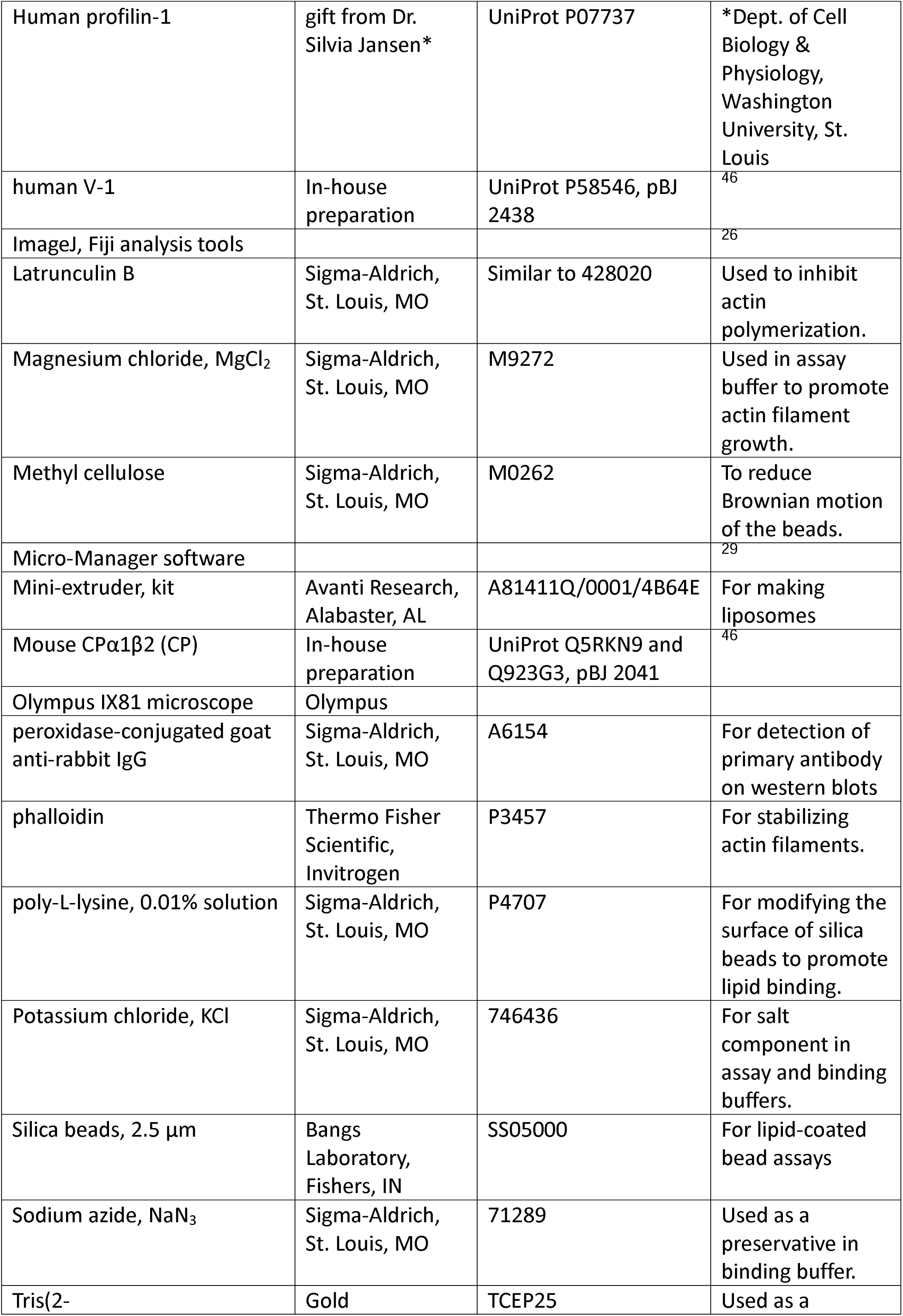
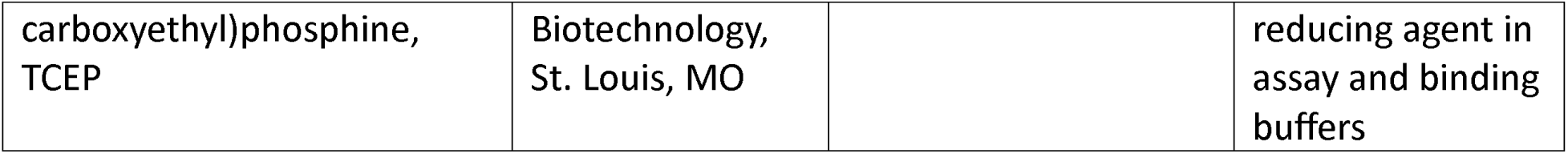

| Name | Company | Catalog Number | Comments |
|---|---|---|---|
| 0.2 µm centrifugal filter | Thermo Fisher Scientific | 50-197-9571 | For collecting beads for SDS-PAGE and WB analysis |
| 0.2 µm polycarbonate membrane | Avanti Research, Alabaster, AL | 610006 | For making liposomes |
| 1,2-dioleoyl-sn-glycero-3-[(N-(5-amino-1-carboxypentyl)iminodiacetic acid)succinyl] (nickel salt) (18:1 DGS-NTA(Ni)) | Avanti Research, Alabaster, AL | 790404 | A Ni-functionalized lipid for coupling His-tagged proteins to the lipid-bead surface |
| 1,2-dioleoyl-sn-glycero-3-phospho-L-serine (sodium salt) (18:1 PS, DOPS) | Avanti Research, Alabaster, AL | 840035C | For making liposomes |
| 1,2-dioleoyl-sn-glycero-3-phosphocholine (18:1 (Δ9-Cis) PC, DOPC) | Avanti Research, Alabaster, AL | 850375C | For making liposomes |
| 1,2-dioleoyl-sn-glycero-3-phosphocholine-N-(Cyanine 5) (18:1 Cyanine 5 PC) | Avanti Research, Alabaster, AL | 850483 | For Cy5 fluorescence to observe the lipid on the bead surface |
| 1% 1,2-dioleoyl-sn-glycero-3-phosphoethanolamine-N-(1-pyrenesulfonyl)(18:1 pyrene PE) | Avanti Research, Alabaster, AL | 810331 | For pyrene fluorescence to observe the lipid on the bead surface |
| (3-Aminopropyl)triethoxysilane, APTES | Sigma-Aldrich, St. Louis, MO | 281778 | For modifying the surface of silica beads to promote lipid binding. |
| 3-(N-morpholino)propanesulfonic acid, MOPS | Sigma-Aldrich, St. Louis, MO | M1254 | For making binding buffer. |
| 4-(2-hydroxyethyl)-1-piperazineethanesulfonic acid, HEPES | Sigma-Aldrich, St. Louis, MO | RDD002 | For making assay buffer. |
| Adenosine triphosphate, ATP | Gold Biotechnology, St. Louis, MO | A-081-25 | For binding to G-actin so it can polymerize. |
| Alexa 647-labeled donkey anti-mouse IgG | Thermo Fisher Scientific | A31571 | For detection of proteins on the bead or within the actin network |
| Alexa 647-labeled goat anti-rabbit IgG | Thermo Fisher Scientific | A21244 | For detection of proteins on the bead or within the actin network |
| Alexa-488 or Alexa-594 G-actin | In-house preparation | UniProt P68135 | <sup>19</sup> |
| Alexa-594 CBR126 | In-house preparation | UniProt Q5VZK9 | <sup>18</sup> |
| Alexa-594 CP | In-house preparation | UniProt Q5RKN9 and Q923G3 but mutations to remove intrinsic cysteines, pBJ 2478 | Rationale and preparation methods as described<sup>45</sup> |
| Alexafluor 594 C5 maleimide | Thermo Fisher Scientific | A10256 | For fluorescent labeling of proteins via a maleimide-cysteine reaction. |
| anti-Arp3 antibody (clone FMS338) | Sigma-Aldrich, St. Louis, MO | A5979 | For detection of proteins on the bead or within the actin network |
| anti-CARMIL1 polyclonal antibody | Thermo Fisher Scientific | 27133-1-AP | For western blot detection of CARMIL1 CBR126 amounts attached to beads or for detection of proteins on the bead or within the actin network |
| anti-N-WASP polyclonal antibody | Thermo Fisher Scientific | PA5-52198 | For western blot detection of VVCA amounts attached to beads |
| Arp2/3 complex, porcine brain | Cytoskeleton Inc. | RP01P | Used to create branched actin networks. |
| b-mercaptoehanol, bME | Sigma-Aldrich, St. Louis, MO | M3148 | To reduce photobleaching in bead assay samples. |
| Bovine serum albumin, BSA | Sigma-Aldrich, St. Louis, MO | A9647 | To prevent non-specific sticking to beads and to glass surfaces. |
| Chemiluminescence substrate | Thermo Fisher Scientific | 32209 | For detection of antibodies on western blots |
| EM-CCD camera | Hamamatsu | C9100 | For imaging bead samples. |
| Ethylene glycol-bis(b-aminoethyl ether)-N,N’,N’-tetraacetic acid, EGTA | Sigma-Aldrich, St. Louis, MO | E3889 | Used as a chelating agent in assay buffer. |
| G-actin from skeletal muscle α-actin | In-house preparation | UniProt P68135 | <sup>19</sup> |
| glutathione-functionalized polystyrene beads | Spherotech, Lake Forest, IL | GSHP-20-5 | For coupling GST-fusion proteins to the beads |
| GST- or His-tagged human N-WASP | In-house preparation | UniProt O00401; fragment P392 to D505 (pBJ 2497 and 2534, WH2-WH2-CA, aka “VVCA”) and fragment P277 to D505 (pBJ 2541, PRD-WH2-WH2-CA, aka “PVVCA” | <sup>18</sup> |
| GST- or His-tagged or untagged human CARMIL1 CBR | In-house preparation | UniProt Q5VZK9; fragment E964 to S1078 (aka “CBR115”, pBJ 2411) and fragment E964 to T1089 (aka “CBR126” pBJ 2491) | <sup>46</sup> |
| Hexa His tag peptide | APExBIO | A6006 | Used as a placeholder to keep the molar His-tag constant on Ni-lipid beads. |
| Human profilin-1 | gift from Dr. Silvia Jansen* | UniProt P07737 | *Dept. of Cell Biology & Physiology, Washington University, St. Louis |
| human V-1 | In-house preparation | UniProt P58546, pBJ 2438 | <sup>46</sup> |
| ImageJ, Fiji analysis tools |  |  | <sup>26</sup> |
| Latrunculin B | Sigma-Aldrich, St. Louis, MO | Similar to 428020 | Used to inhibit actin polymerization. |
| Magnesium chloride, MgCl<sub>2</sub> | Sigma-Aldrich, St. Louis, MO | M9272 | Used in assay buffer to promote actin filament growth. |
| Methyl cellulose | Sigma-Aldrich, St. Louis, MO | M0262 | To reduce Brownian motion of the beads. |
| Micro-Manager software |  |  | <sup>29</sup> |
| Mini-extruder, kit | Avanti Research, Alabaster, AL | A81411Q/0001/4B64E | For making liposomes |
| Mouse CPα1β2 (CP) | In-house preparation | UniProt Q5RKN9 and Q923G3, pBJ 2041 | <sup>46</sup> |
| Olympus IX81 microscope | Olympus |  |  |
| peroxidase-conjugated goat anti-rabbit IgG | Sigma-Aldrich, St. Louis, MO | A6154 | For detection of primary antibody on western blots |
| phalloidin | Thermo Fisher Scientific, Invitrogen | P3457 | For stabilizing actin filaments. |
| poly-L-lysine, 0.01% solution | Sigma-Aldrich, St. Louis, MO | P4707 | For modifying the surface of silica beads to promote lipid binding. |
| Potassium chloride, KCl | Sigma-Aldrich, St. Louis, MO | 746436 | For salt component in assay and binding buffers. |
| Silica beads, 2.5 µm | Bangs Laboratory, Fishers, IN | SS05000 | For lipid-coated bead assays |
| Sodium azide, NaN<sub>3</sub> | Sigma-Aldrich, St. Louis, MO | 71289 | Used as a preservative in binding buffer. |
| Tris(2-carboxyethyl)phosphine, TCEP | Gold Biotechnology, St. Louis, MO | TCEP25 | Used as a reducing agent in assay and binding buffers |

